# Characterising genome architectures using Genome Decomposition Analysis

**DOI:** 10.1101/2021.12.01.470736

**Authors:** Eerik Aunin, Matthew Berriman, Adam James Reid

**Affiliations:** Wellcome Sanger Institute, Cambridge, CB10 1SA, United Kingdom; The Gurdon Institute, University of Cambridge, Cambridge CB2 1QN, United Kingdom

**Keywords:** Comparative Genomics, Genome Architecture, Apicomplexa, Tree of Life, Software Pipeline, Chromosomes

## Abstract

Genome architecture describes how genes and other features are arranged in genomes. These arrangements reflect the evolutionary pressures on genomes and underlie biological processes such as chromosomal segregation and the regulation of gene expression. We present a new tool called Genome Decomposition Analysis (GDA) that characterises genome architectures and acts as an accessible approach for discovering hidden features of a genome assembly. With the imminent deluge of high quality genome assemblies from projects such as the Darwin Tree of Life and the Earth BioGenome Project, GDA has been designed to facilitate their exploration and the discovery of novel genome biology. We highlight the effectiveness of our approach in characterising the genome architectures of single-celled eukaryotic parasites from the phylum *Apicomplexa* and show that it scales well to large genomes.

**Significance:** Genome sequencing has revealed that there are functionally important arrangements of genes, repetitive elements and regulatory sequences within chromosomes. Identifying these arrangements requires extensive computation and analysis. Furthermore, improvements in genome sequencing technology and the establishment of consortia aiming to sequence all species of eukaryotes mean that there is a need for high throughput methods for discovering new genome biology. Here we present a software pipeline, named GDA, which determines the patterns of genomic features across chromosomes and uses these to characterise genome architecture. We show that it recapitulates the known genome architecture of several Apicomplexan parasites and use it to identify features in a recently sequenced, less well-characterised genome. GDA scales well to large genomes and is freely available.

## Introduction

Genome architecture is the arrangement of functional elements within the genome (Koonin 2009) and can be thought of in a linear fashion, or in the three-dimensional arrangement found in nuclei (Rowley & Corces 2018). The architecture of genomes differs greatly across the tree of life. For example, bacteria tend to have small genomes, consisting mainly of single-exon protein coding genes, often arranged in coexpressed operons, with well-defined regulatory regions (Koonin 2009). Eukaryotic genomes are diverse, ranging from those that are relatively compact, with genes lacking introns (e.g. *Leishmania* spp.), to large, repeat-rich genomes, sparsely populated by multi-exon genes with large introns which employ long range regulatory interactions (Lynch & Conery 2003). Although we have an excellent understanding of the evolution of protein-coding genes and how they are shaped by natural selection, we know very little of the forces that shape many aspects of genome architecture, and random drift may be the dominant force in many eukaryotic genomes (Lynch et al. 2011). Despite this, there are many features of genome architecture that are functional and which provide clues to understanding more about the biology of an organism and its evolutionary history. For instance, in the parasitic protozoan *Plasmodium falciparum*, genes involved in evading host immunity are located in the subtelomeric regions of chromosomes where the heterochromatic environment enables clonal variability in gene expression (Lopez-Rubio et al. 2009; Flueck et al. 2009). In mammals, the immunoglobulin and T-cell receptor loci comprise ordered arrays of duplicated genes, allowing the generation of variant antibody and T-cell receptor proteins (Tonegawa 1983). Operons of co-expressed genes are found in some eukaryotes such as kinetoplastids (Johnson et al. 1987) and nematodes (Spieth et al. 1993). Some fungi have genomes in which different regions have distinct evolutionary rates (https://www.sciencedirect.com/science/article/pii/S1749461320300257?via%3Dihub). There are also chromosomes that have distinct architectural patterns within a genome. These include sex chromosomes (C. elegans Sequencing Consortium 1998) and accessory B chromosomes, such as those found in plants and fungi (Ahmad & Martins 2019). In the nematode worm *C. elegans*, repetitive sequences have accumulated mostly at the ends of chromosomes (C. elegans Sequencing Consortium 1998). However, some repeat families have their own distinctive patterns that are repeated across each chromosome, suggesting a variety of forces at work (Surzycki & Belknap 2000).

A key problem hampering our understanding of genome architecture has been a lack of chromosome-scale genome assemblies. However, steady advancements in the quality of long-read genome sequencing (Wenger et al. 2019) and scaffolding technologies (Burton et al. 2013; Kaplan & Dekker 2013) are beginning to solve this. Furthermore, projects such as the Darwin Tree of Life (https://www.darwintreeoflife.org/) and the Earth BioGenome Project (https://www.earthbiogenome.org/) are planning to deliver chromosome-scale assemblies for all species across the eukaryotic kingdom. A second problem is that there is no recognised approach for characterising chromosome architectures, something that would greatly facilitate studies on their evolution.

We present a new approach to characterise the linear architecture of genomes called Genome Decomposition Analysis (GDA). A genome sequence is divided into windows of arbitrary length and features are calculated for each window. Features can be derived solely from the sequence itself, including GC content, protein-coding potential, and repeat content, or include properties derived from other sources, such as sequence homology, gene expression, chromatin modifications, and recombination frequencies. The dimensionality of the resulting data matrix of windows and features is reduced and the results clustered. Parameters are explored to produce distinct clusters with a minimum of unclassified windows. Features are then identified that characterise these clusters. The pattern of clusters across chromosomes is inspected to reveal, for example, that the middles of chromosomes are distinct from the ends and that they are enriched in repeats. GDA includes an easy-to-use web application for data exploration and visualisation.

Apicomplexan parasites are well-studied due to their importance in disease and have well-understood genome architectures, making them ideal candidates for developing and testing GDA. We use GDA to: (i) refine our earlier definition of the genome architecture of the malaria parasite *P. falciparum* and characterise variation in its relatives; (ii) show that bands of repeat-rich sequence cover all chromosomes of the chicken parasite *Eimeria tenella* and compare its architecture to that of the canonical coccidian *Toxoplasma gondii*, revealing they both have distinctive but gene-poor subtelomeres; and (iii) demonstrate the potential of GDA for understanding the genome architecture of much larger genomes.

GDA is under the MIT licence and is available from GitHub: https://github.com/eeaunin/gda

## Results

### Design of the GDA pipeline

We developed GDA to identify features of genome architecture from highly contiguous genome assemblies as a basis for further study of genome evolution. The tool consists of three main parts: a genomic feature extraction pipeline that calculates feature values in windows across the genome; dimension reduction and clustering of these windows; and visualisation and data exploration using a web-browser application (Figure 1). The minimal required input for the pipeline is a genome assembly FASTA file. The features that are extracted from the FASTA file are: GC content, GC skew, AT skew, CpG dinucleotide frequency, *k-*mer frequencies, stop codon frequency, matches to a telomeric sequence motif, low complexity sequence content, tandem repeat content, coverage of simulated reads, retrotransposons, inverted repeats and repeat families (Supplementary Table 1). A more exhaustive repeat analysis can be included by running RepeatModeler which produces features describing the distribution of individual complex and simple repeats as well as features describing the sums of complex and simple repeats. Gene annotations can be used to produce bedgraph tracks of mRNA, tRNA and rRNA gene densities, average exon count, exon length and intron length. Where gene annotation files are unavailable, the pipeline can annotate genes. Likewise, if proteome FASTA files are provided for related species, the pipeline can produce bedgraph tracks based on the counts of predicted paralogs, orthologs, conserved proteins and species-specific proteins. It is also possible to add any user-generated tracks, using coordinates of the genome being analysed, to be included as input to the clustering step.

**Figure 1.**
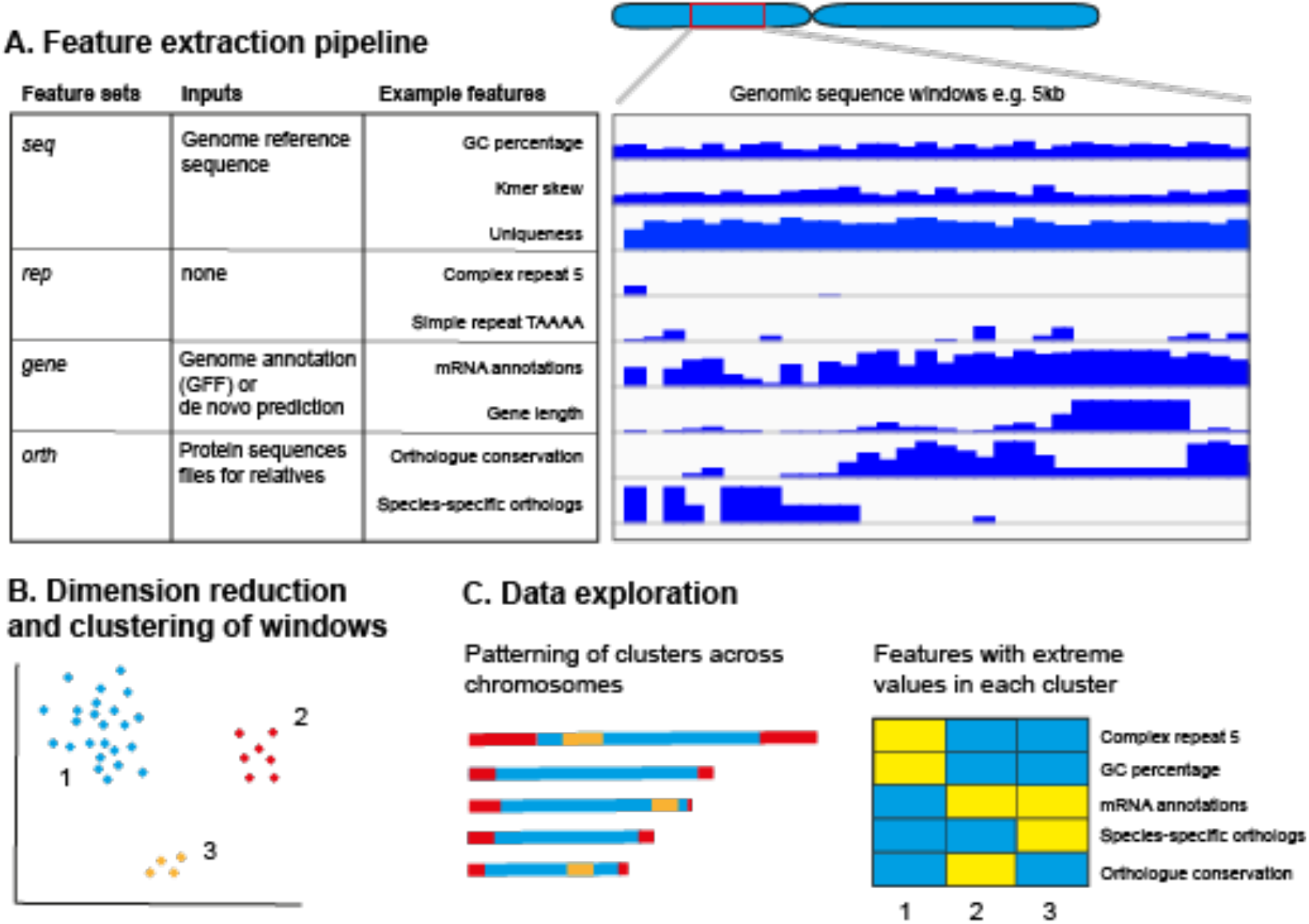
Overview of the GDA pipeline. (A) Feature sets are derived from the genome reference sequence (*seq*), repeat finding (*rep*), gene annotations (*gene*) and evolutionary relationships between genes (*orth*). The genome is divided into user-defined, non-overlapping windows (e.g. 5kb in length) from which the value of each feature is determined. (B) The resulting matrix of feature values per window is embedded in two dimensions and clustered to identify groups of windows with similar properties. (C) The data can be explored in a number of ways using a web-browser based app. The clustering labels are mapped back to the chromosomes to highlight architectural features and a heatmap displays the features which define the clusters.

Each feature is examined in sliding windows across the genome, the output of which is stored in bedgraph files. These bedgraph files can be visualised in a genome browser such as IGV (Robinson et al. 2011). The data in the bedgraph files are merged into a tab separated (TSV) file and are scaled to fit the range between 0 and 1. The resulting table is then analysed using UMAP, a dimensionality reduction approach (McInnes et al. 2018). HDBSCAN (McInnes et al. 2017) is then run to detect clusters of genomic windows in the UMAP embedding. Next, the user can explore different values of key parameters for UMAP and HDBSCAN and compare the clusterings obtained. When suitable parameter values have been chosen, the clustering and analysis script is run, producing a set of output files. One of the output files is a BED file that marks which cluster each genomic window belongs to. We identify characteristic features for each cluster using the Kolmogorov-Smirnov test. The clustering and analysis results can be explored using the GDA web app that includes a scatter plot of clustered windows, how these clusters are arranged over the genome, heatmaps of features enriched in each cluster, and the cluster composition of scaffolds.

### Redefining *Plasmodium falciparum* genome architecture

A complete chromosomal genome assembly of the human malaria parasite *Plasmodium falciparum* has been available for almost 20 years (Gardner et al. 2002; Böhme et al. 2019). Given the importance of the *P. falciparum* genome as a reference for studying one of the most persistent and deadly human infectious diseases, it is not surprising that there is a good understanding of its architecture. More surprising is that it has been difficult to formally define what is known intuitively from having studied it.

We first tested the ability of GDA to identify the known architectural features of the *P. falciparum* genome using only features derived from the genome sequence itself (*seq* feature set). We chose a window size of 5kb to capture a small number of genes per window and to reflect the resolution of the genome architecture we expect to see. For genomes where the architecture is unknown we recommend choosing several window sizes and comparing results. We explored a range of UMAP *nearest neighbour* (n) and HDBSCAN *minimum cluster size* (c) parameters but picked n = 5 and c = 50 as these resulted in a relatively high silhouette score of 0.28, with 100% of windows being classified (Sup Fig 1; Figure 2A). The three resulting clusters defined the core (cluster 2), the multigene family arrays (cluster 1) and the GC-rich Telomere Associated Repeat Element (TARE) region adjacent to the telomeres (cluster 0; Figure 2B). The core was characterised by uniqueness of sequence (simulated mapping coverage of 9.94x, p= 1.23e-96), tandem repeats (p= 1.09e-36) and low GC percentage (18.6% vs. 22.2%, p= 8.43e-43) (Figure 2C). The multigene family-rich regions were defined by high CpG percentage (0.96 vs. 0.66, p= 9.00e-32) and low uniqueness as measured by mapping coverage of simulated reads (3.9x compared to expected 10x, p= 4.44e-131). This was caused by highly similar regions in tandemly duplicated gene clusters. The TARE region was defined by high GC percentage (32.4%, KS test p-value = 4.90e-146), high stop codon frequency (0.24, KS test p-value 1.43e-87), and *k*-mer deviation (3-mer, p=1.30e-69 and 4-mer, p= 6.69e-48) (Figure 2C). This definition of *P. falciparum* genome architecture required only the genome sequence and simple parameters derived from it, yet characterised both the relatively GC-rich telomere-adjacent regions, gene-family rich subtelomeres and the conserved core.

**Figure 2.**
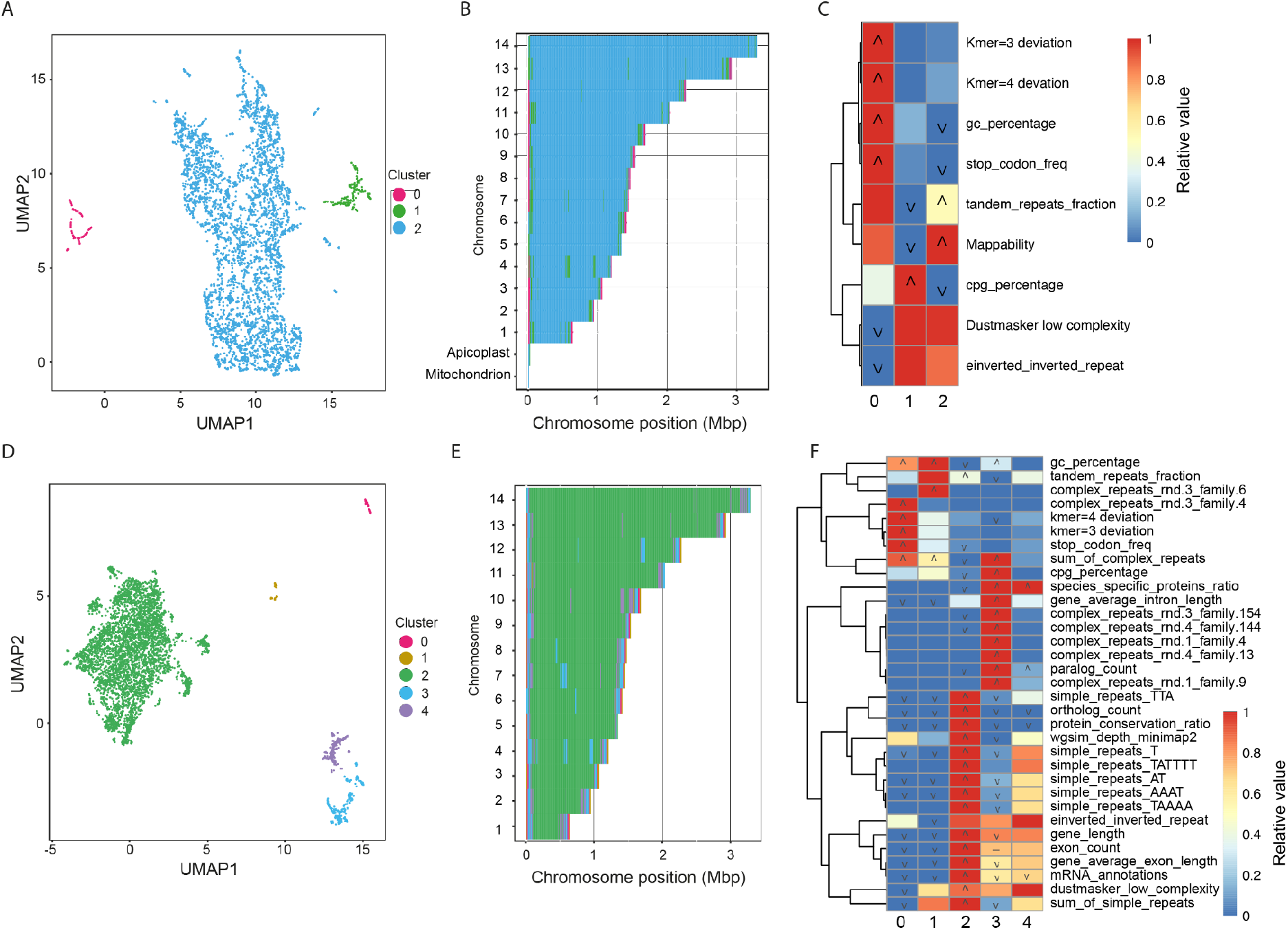
GDA analysis of the *Plasmodium falciparum* genome. (A) UMAPembedding (n = 5) and HDBSCAN2 clustering (c = 50) of 5kb windows using simple features derived from the genome sequence (seq feature set). (B) Projection of clusters onto the chromosomes highlights the localisation of cluster 0 windows at the very ends of chromosomes, with cluster 1 windows adjacent to these and within the cores of some chromosomes. (C) Heatmap showing features enriched in each cluster with seq feature set. Colours indicate the relative value of the feature in each cluster (red = highest, blue lowest), icons indicate significance (‘∧’ = KS test greater p-value <= 1e-20, ‘∨’ = KS test lesser p-value <= 1e-20, ‘-’ = great and lesser p-values <= 1e-20) (D) UMAP embedding (n = 20) and HDBSCAN2 clustering (c = 50) of 5kb windows with seq+gene+rep+orth feature set. (E) Projection of clusters onto chromosomes shows that the additional features break the subtelomeric regions into four distinct regions and that two types of island (clusters 3 and 4) interrupt the core (cluster 2) on some chromosomes. (F) Heatmap showing features enriched in each cluster with all features.

To improve on this definition of the genome architecture we generated features from three additional sources, adding gene annotations (*seq+genes*), then repeat classification (*seq+genes+reps*) and finally protein-coding gene conservation (*seq+genes+reps+orths*). For each of these feature sets we re-ran the feature extraction pipeline and chose clustering parameters that minimised the number of unclustered windows, while providing a number of large well-separated clusters, with a good silhouette score. Adding gene annotations altered the definition of the multigene family-rich subtelomeres, including the smaller, more well-conserved families, because both these regions and those containing high copy number multigene families are less gene-dense than the core (Figure 3). Adding repeat classification (*seq+genes+reps*) differentiated the TARE2-5/SB-2 region (named *complex_repeats_rnd-3_family-6* by GDA) closest to the telomeres (Gardner et al. 2002) from the TARE6/SB-3/rep20 repeat (named *complex_repeats_rnd-3_family-4* by GDA). Repeat identification altered the definition of the multigene family regions to be more like that found when only sequence-based information was used. This was because the larger multigene families were identified as repeats and this excluded the smaller multigene families. Including all this information, plus analysis of gene conservation (*seq+genes+reps+orths*) allowed improved definition of the large multigene family-containing subtelomeric cluster - all 65 *var* genes, 155/157 rifin genes and 31/32 *stevor* genes overlapped cluster 3. It also highlighted the more conserved, distal subtelomeric regions containing smaller gene families, where there is conservation of synteny within *P. falciparum*, but not between species (cluster 4; Figure 2E; Figure 3B).

**Figure 3.**
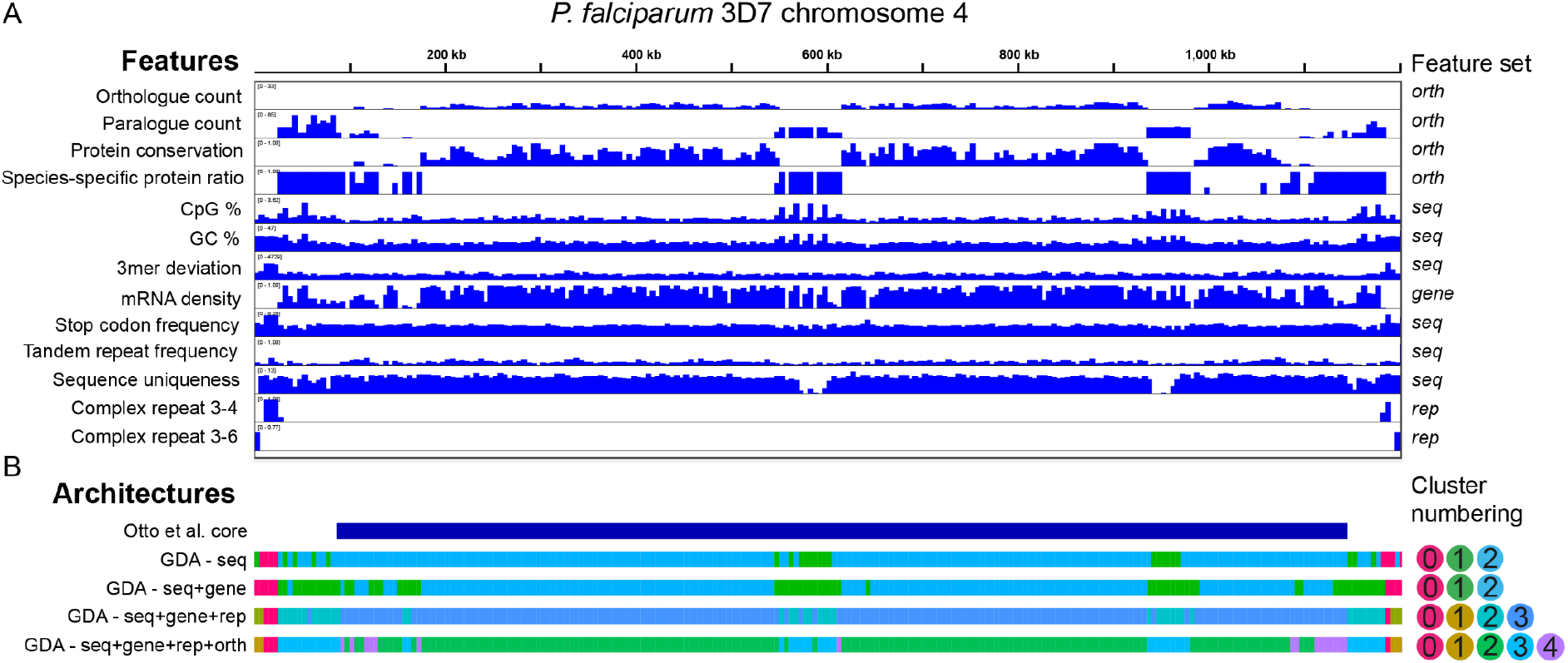
Detailed view of *Plasmodium falciparum* chromosome 4. (A) A selection of the features used as input to GDA displayed across the 1.2Mb chromosome 4. These features were identified as significant in one or more clusters of one or more GDA runs. (B) Definition of chromosome architectures based on Otto *et al*. (Otto et al. 2018). GDA was run with basic sequence features, with the addition of gene annotation, with gene annotations and complex repeat finding, with gene annotations, complex repeat finding and orthology analysis.

### Defining the unique arrangement of the *P. knowlesi* genome

Most *Plasmodium* species have similar genome architectures to *P. falciparum*. The clear exception is *P. knowlesi*, a related species that also causes malaria in humans and other primates. In this species, the largest, most rapidly evolving multigene families (in this case *sicavar* and *pir*) are found in islands throughout chromosomes associated with telomere-like repeats (Pain et al. 2008). We used this example to examine the utility of GDA for comparative genomics - identifying differences in architecture between related species. *P. vivax* is a closer relative to *P. knowlesi* than *P. falciparum* but has a genome split into gene-family rich subtelomeric regions and a well conserved core like *P. falciparum*. We ran GDA on the *P. vivax* genome with a *seq+gene+rep+orth* feature set, identifying two clusters characterising the whole genome. This confirmed that like *P. falciparum*, most of *P. vivax* chromosomes are made up of cores with well-conserved genes (cluster 0; Figure 4A-C). Conversely, the subtelomeres contain species-specific genes with high numbers of paralogues (cluster 1).

**Figure 4.**
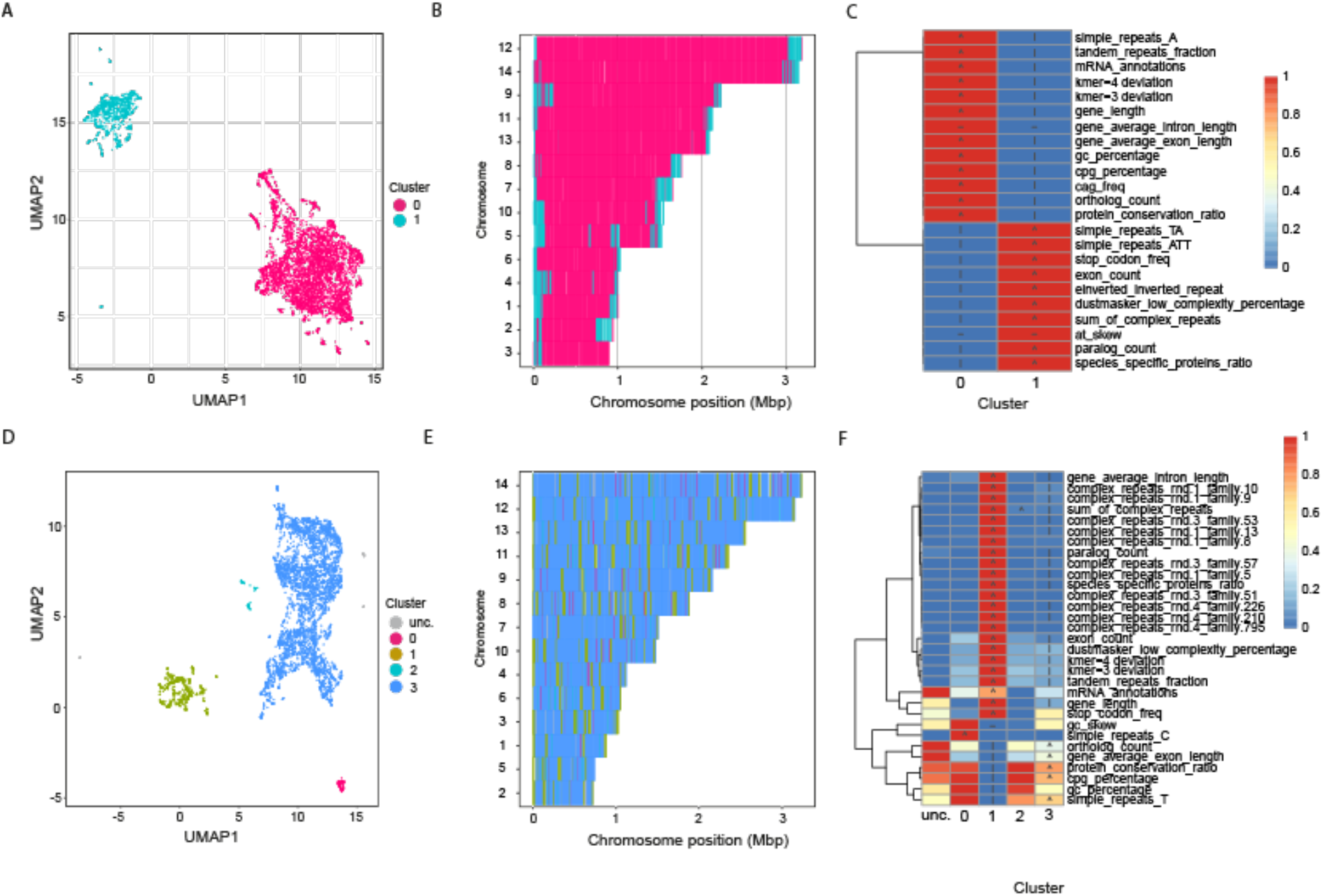
GDA analysis of the *Plasmodium vivax* P01 and *P. knowlesi* H genomes. (A) The *P. vivax* genome neatly separates into two clusters with *seq+rep+gene+orth* feature sets. (B) These represent core (magenta) and subtelomeric (cyan) regions, which are typified, amongst other things, by having one-to-one orthologous genes versus highly paralogous species-specific genes, respectively (C). (D) *P. knowlesi* separated into four clusters, with no clear subtelomeric localisation (E). (F) The cluster with large species-specific gene families equivalent to the subtelomeric cluster of *P. vivax* (cluster 1; green) is dispersed throughout each chromosome.

GDA analysis of *P. knowlesi* resulted in four clusters with 82.96% of the windows assigned falling into cluster 3, representing well-conserved genes. Cluster 1 (12.41%) represented the multi-gene family-rich regions which are interspersed throughout the chromosomes, rather than concentrated towards the telomeres as observed in other *Plasmodium* spp. This cluster was also enriched for complex repeat families (*sum of complex repeats* p=0). Several of these repeat families contained telomere-like repeats (e.g. TT[T/C]AGGG) as expected from previous analysis (Pain et al. 2008). Cluster 2 made up 1.8% of the genome and was enriched only for *simple_repeats_C* (p= 1.47e-176). This relates to a previously unidentified feature of the genome: 63 polyC repeats of ∼20 nucleotides. Twenty-eight of these repeats were found in introns, while others tended to lie close to genes. Here, GDA makes clear the alteration in genome architecture between closely related species, while also identifying previously hidden features.

### Identification of repeat-rich bands and large gene-poor subtelomeres in *Eimeria tenella*

*Eimeria spp.* parasites have been found in a wide range of vertebrates and commonly cause coccidiosis in domesticated chickens. We have previously shown that their ∼50 Mb genomes contain a banded pattern of regions rich in CAG and telomere-like (TTTAGGG) repeats (Reid et al. 2014). Coding regions are enriched for the CAG repeat, which tends to encode Homopolymeric Amino Acid Repeats (HAARs) of alanine, serine or glutamine and litter even very well-conserved genes. We recently sequenced the genome of *Eimeria tenella* using long reads and Hi-C scaffolding, producing a nearly chromosomal assembly (Aunin et al. 2021).

We investigated whether GDA was able to identify the repeat rich bands and other distinctive features in this genome using the new chromosome-scale assembly. Using the *seq* feature set resulted in three clusters. Figure 5 shows that this simple input was sufficient to define the repeat-rich bands across the genome. 95.2% (3,927) of genes containing HAARs fell into cluster 1. This highlights that when using only simple features, GDA is able to accurately capture this aspect of genome architecture, and furthermore, that *E. tenella* genome architecture is dominated by this feature.

**Figure 5.**
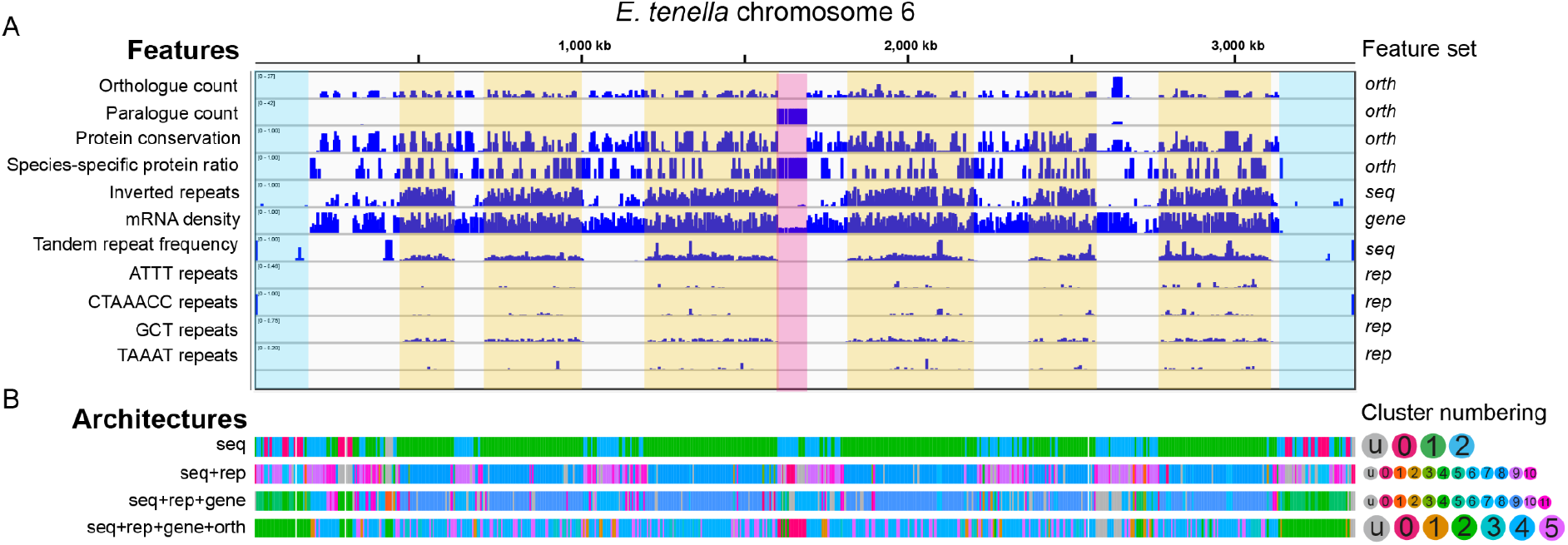
Repeat-rich bands and gene-poor subtelomeres of *Eimeria tenella* are captured more or less well by different feature sets. (A) A number of features are shown in 5kb windows across chromosome 6 of *E. tenella*. The repeat-rich bands, defined here by GCT (CAG) repeats are highlighted in yellow. The gene-poor subtelomeres are highlighted in blue and a *sag* multigene family array in pink. (B) Four different architectures, based on different feature sets are shown below. The *seq*, *seq+rep* and *seq+rep+genes* feature sets capture the repeat-rich regions very well, with the last of these also capturing the gene-poor subtelomeres. The *seq+rep+gene+orth* feature set does not capture the repeat-rich regions in a single cluster but instead focuses more on whether a window contains more well-conserved genes or not. It retains the cluster identifying the gene-poor subtelomeres and highlights arrays of *sag* genes.

To better understand the repeats present in the different regions, we ran GDA again, adding in the *rep* feature set (silhouette score = 0.32; Figure 6A). We saw that cluster 8 (41.69% of the genome) was enriched for simple_repeats_CTAAACC (p = 0; i.e. the telomere-like repeat) and simple_repeats_GCT (p = 0; i.e. CAG repeat) as well as inverted repeats and several complex repeat families (Figure 6C). This cluster overlapped 93.8% of HAARs (26,728/28,483). With this feature set, cluster 9 represented the gene-rich parts of the genome lacking repeats (23.61%), while cluster 10 (9.04%) —intermediate between clusters 8 and 9 in the UMAP plot — was enriched in inverted repeats and sum of complex repeats. Cluster 5 captured the LTR retrotransposons, which are not a common feature in apicomplexan genomes and were first identified in *E. tenella* and then subsequently in avian malaria parasites (Ling et al. 2007; Böhme et al. 2018). Cluster 4 was enriched for TGTTGC repeats, which were the only enriched simple repeats to not colocalise in the repeat-rich cluster 8 regions, instead being more evenly dispersed throughout the chromosomes. On chromosome 6 it is repeated between tRNA genes in a tRNA cluster, but otherwise does not have an obvious pattern.

**Figure 6.**
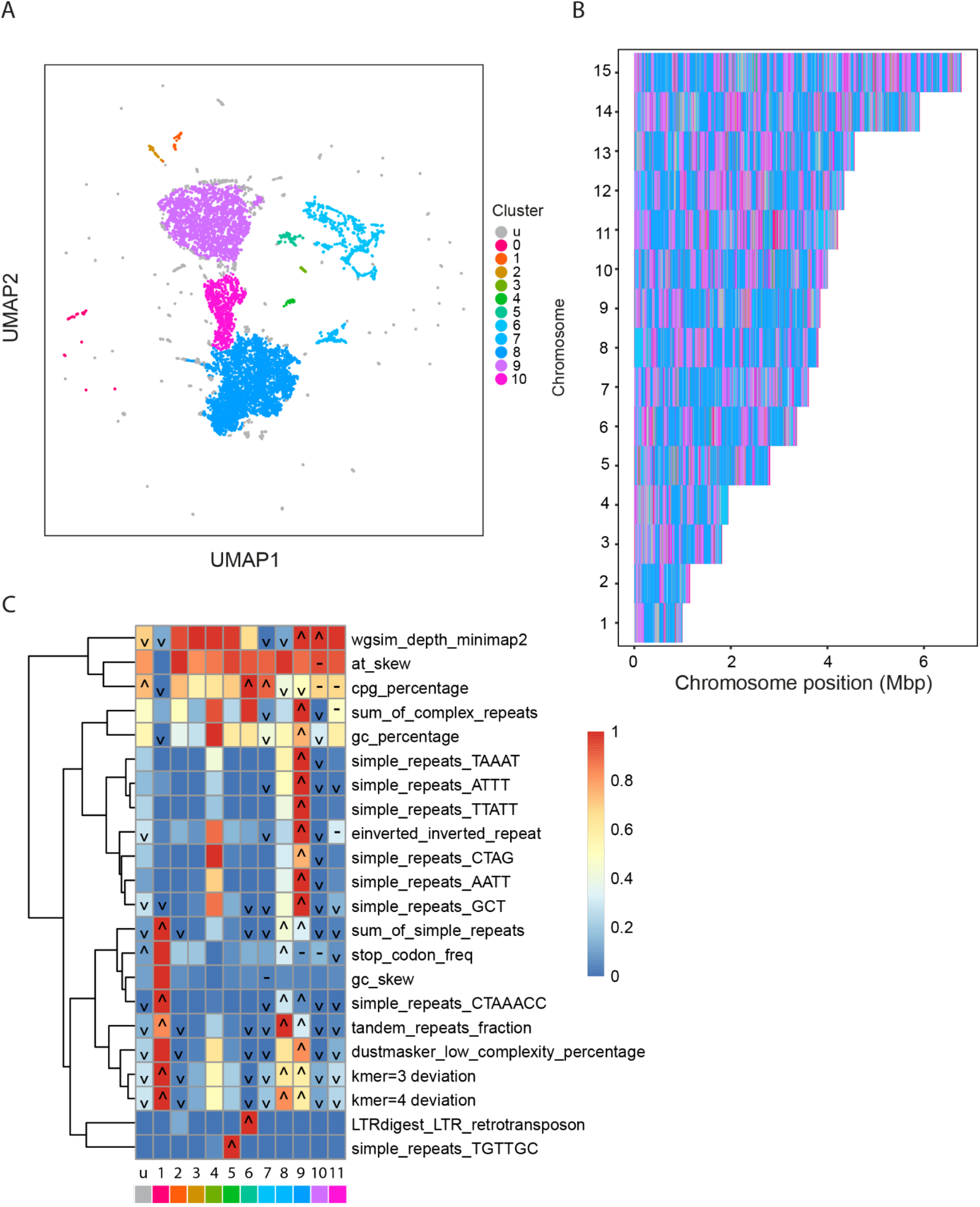
GDA analysis of *Eimeria tenella* with the *seq+rep* feature set. (A) Analysis of *E. tenella* with the *seq+rep* feature set identified 11 clusters. The majority of the genome was separated into three or four clusters found in bands across each chromosome (B). (C) These include the repeat rich region (cluster 8; dark blue), a cluster which is similar but lacks repeats (9; purple) and an intermediate cluster (10; magenta) which is enriched for *sum of complex repeats* and *inverted repeats*, but not the GCT/CAG and telomere-like (CTAAACC) repeats found in cluster 8.

Adding in gene features (*seqs+reps+genes*) distinguished gene-poor regions at the subtelomeres and internally within chromosomes (Figure 5). Clusterings with high silhouette scores and relatively few unclassified windows failed to distinguish the repeat-rich regions. We picked parameters which resulted in a separate cluster for windows intermediate between repeat-rich and repeat-poor clusters, with a relatively moderate 13.68% unclassified windows and silhouette score = 0.18 (n=10, c=50; Sup Fig 2). This allowed the identification of gene-poor subtelomeric (and sometimes internal) regions with repeat-rich regions still well-characterised (26,566/28,483 HAARs in cluster 9; Figure 5). Gene-poor subtelomeric regions have not previously been described as a feature of *Eimeria* chromosomes. These subtelomeric gene deserts (clusters 4 and 5) have high CpG content and cluster 5 has high stop codon frequency, while cluster 4 has low uniqueness, despite not being enriched for any particular repeat families.

We wanted to determine whether gene poor subtelomeres were also present in other *Coccidia* and so we ran GDA on the related species *Toxoplasma gondii* with seq+gene+rep+orth feature sets. The genome resolved into 5 distinct clusters, with no unclassified windows (Figure 7). Chromosomes often ended in gene-poor regions falling into cluster 1 (mRNA_annotations lower than other regions; p=5.6e-310). These had high stop codon frequency (p=5.35e-74), high GC skew (p=8.05e-30) and were enriched for complex repeats (p = 1.03e-26), although no individual repeats in particular, much like *E. tenella*.

**Figure 7.**
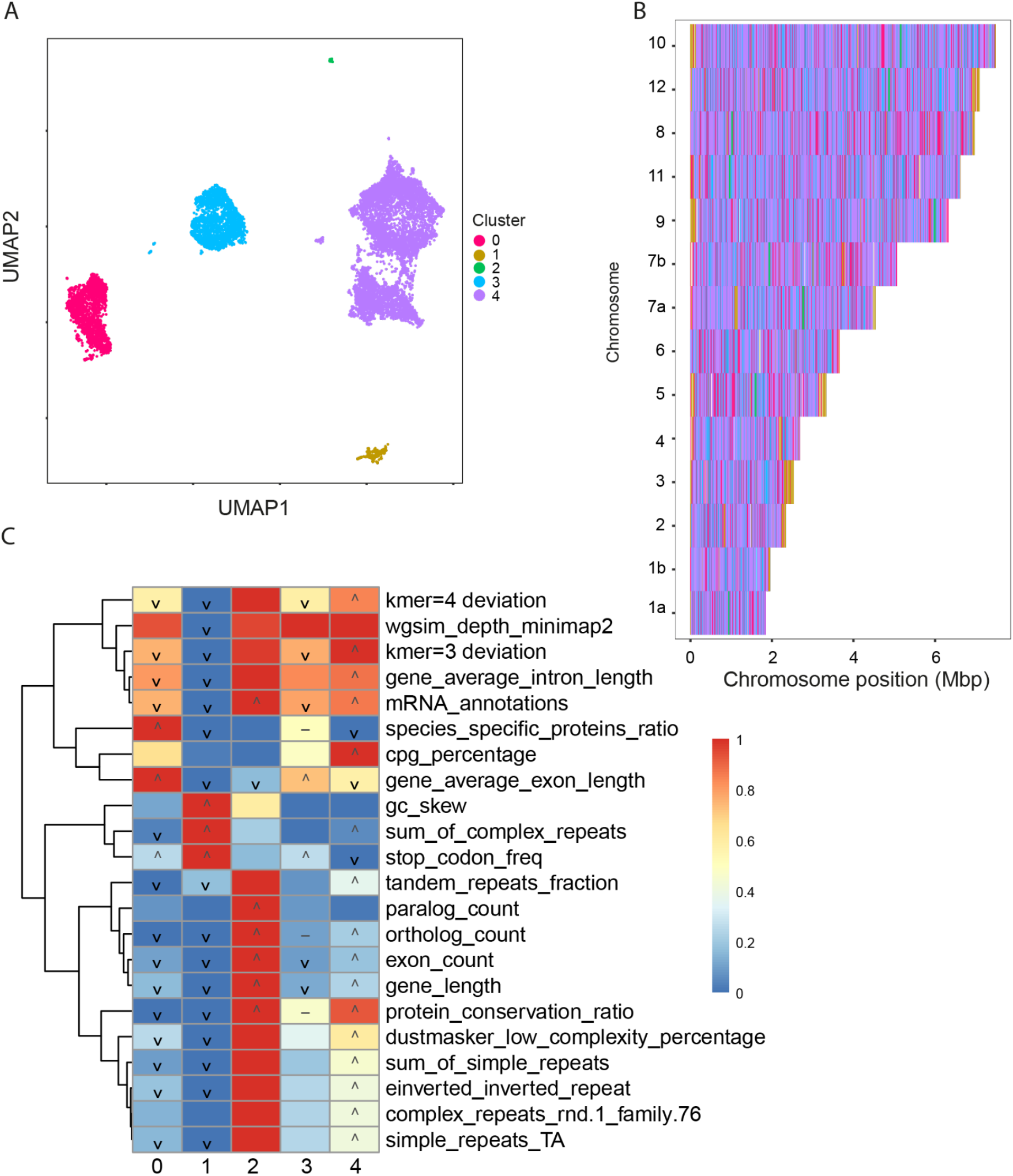
GDA analysis of *Toxoplasma gondii* highlights gene-poor subtelomeres and gene family-rich islands. (A) Using the *seq+rep+gene+orth* feature set, the *T. gondii* genome separated into 5 distinct clusters. (B) Cluster 1 (gold) was often found at the ends of chromosomes and was typified by low numbers of mRNA annotations, high GC skew, complex repeats and stop codon frequency (C). This is similar to what we see in *E. tenella* subtelomeres.

Next we ran GDA on *E. tenella* including the *orth* feature set (*seqs+reps+genes+orths*) to see if we could identify patterns of gene conservation amongst the complexity of the repeat regions (6 clusters, n=10, c=100, 1.9% windows unclassified). The gene-poor subtelomeres remained well classified (Figure 5), but the *sag* gene arrays on chromosomes 6, 9 and 11 were now also well-captured by cluster 0. Of 78 *sag* genes, 51 overlapped cluster 0 windows. In this clustering the repeat-rich cluster was lost. Instead, much of each chromosome was split into windows with well-conserved genes (cluster 4 – 46.61% windows) or more species-specific genes (cluster 5 – 15.61%). Both these clusters were enriched for “simple_repeat_GCT” (i.e. CAG repeats; KS-test one-sided p-value 1.03e-234 for cluster 4, 1.43e-82 for cluster 5).

The *E. tenella* genome highlights how some important properties of genome architecture are not well captured with a single parameter set. Using different feature sets, and parameters such as window size, enabled different aspects of genome architecture to be represented.

### GDA can be run on large genomes and with high resolution

We measured the time taken to run the genomic feature extraction pipeline of GDA with the genome assemblies of four different species representing a range of genome sizes: *Plasmodium falciparum* (∼23Mb), *Caenorhabditis elegans* (∼100Mb), *Schistosoma mansoni* (∼410Mb) and *Homo sapiens* (∼3300Mb) (Table 1). In each case 5kb windows were used, meaning that for *H. sapiens*, features were calculated over 654,762 windows. Memory requirements were roughly correlated with genome size and were not greatly affected by repeat finding. Run time was roughly correlated with genome size, however *C. elegans* took longer to process than *S. mansoni*. The major factor contributing to long run times was using RepeatModeler to identify repeats *de novo (rep* feature set). Without this step, analysis of the *P. falciparum* genome was completed in 16 minutes and the human genome in less than 11 hours. When *de novo* repeat finding was included these analyses took ∼16 hours and 92 hours respectively. However, it is clear that GDA can be run effectively on large genomes with resources commonly available on bioinformatics compute clusters, even including time-intensive repeat finding.

**Table 1.** Resource requirements for running the GDA feature extraction pipeline on a range of genomes. The GDA feature extraction pipeline was run with four genomes of different sizes. De novo repeat detection had a large effect on run time while genome size caused increases in both run time and memory usage.

| Feature set |  | *P. falciparum* | *C. elegans* | *S. mansoni* | *H. sapiens* |
| --- | --- | --- | --- | --- | --- |
|  | Assembly size (Mb) | 23.33 | 100.29 | 409.57 | 3,272.09 |
| *Seq+gene+orth*<br>(without RepeatModeler) | Run time | 16 min | 2 h 56 min | 1 h 57 min | 10 h 30 min |
|  | CPU time (s) | 2,345 | 35,288 | 17,410 | 122,967 |
|  | Max memory use (Mb) | 4,454 | 11,798 | 12,042 | 129,378 |
| *Seq+gene+rep+orth* (with RepeatModeler) | Run time | 15 h 51 min | 19 h 2 min | 44 h 56 min | 92 h 12 min |
|  | CPU time (s) | 573,551 | 479,531 | 1,265,211 | 1,916,769 |
|  | Max memory use (Mb) | 4,379 | 11,218 | 12,622 | 135,828 |

## Discussion

We have presented a new tool, GDA, which decomposes a genome sequence into windows, identifying those with similar properties and enabling the characterisation of genomic architectural features. This is achieved most simply using properties derived from the genome sequence alone, but a wide range of additional properties can be used as input. We have shown that GDA recapitulates the well-described architecture of the malaria parasite *Plasmodium falciparum* and in doing so defines regions of interest that can be further explored. The description of the *P. falciparum* genome was robust to different feature sets, suggesting that each part of the genome has multiple features distinguishing it from other regions which are correlated with each other. In the *Eimeria tenella* genome, GDA analysis highlighted the banded pattern of repeats observed previously (Reid et al. 2014; Ling et al. 2007) and shows for the first time that it is present across all chromosomes. A previous attempt to define these regions involved arbitrary cutoffs, but GDA provides a straightforward and data-driven approach to define the repeat-rich regions. This will facilitate the comparison of different *Eimeria* spp. genomes in studying the evolution of these repeat-rich regions across species.

The power of GDA lies in the way it allows visualisation of genome architecture to suggest hypotheses about genome function and evolution. Applied to closely related species, substantial changes in organisation of genomic features can be quickly recognised (as in the example of *P. vivax* and *P. knowlesi*). The drivers of these features can be readily determined and investigated (as in the CAG repeats in protein-coding genes of *E. tenella*). This makes GDA a powerful tool for any *de novo* genome sequencing or comparative genomics project involving well-assembled genomes. We foresee a range of applications such as sex and accessory chromosome identification, genome assembly curation and interpretation of epigenemic datasets (e.g. ChIP-seq/ATAC-seq). In fact, similar approaches to ours have been used to analyse patterns of chromatin modifications in isolated genomic regions (Nielsen et al. 2012) and patterns of relatedness across genomes (Li & Ralph 2019). However, we are not aware that similar approaches have been applied to characterise genome-wide architecture and we have not found any tool which has this aim.

When considering application of GDA for different purposes and on different sizes of genome, window size is an important parameter. The choice of window size should reflect the resolution of features that the user is interested in. A window size of 1kb in a 100Mb genome may reflect individual parts of genes such as separate exons, introns and promoters which would be appropriate for understanding patterns in many types of ChIP-seq data. On the other hand, windows of 5-10kb may reflect one or a handful of genes or complex repeats per window, while 1Mb windows will reflect more broad aspects of genome architecture.

All Apicomplexan genomes appear to be relatively small and compact, however their architectures are diverse. Unlike some larger genomes, in which there is little linear architectural coherence based on sequence properties, repeats and homology, these genomes display quite definite ordering. Current work on mammalian genomes suggests that important aspects of architecture relating to the control of gene expression are manifest in the third dimension, i.e. the arrangement of the linear chromosomes in space (Yu & Ren 2017). These arrangements can be assayed by techniques such as Chromatin Conformation Capture (e.g. Hi-C). Although not linear in nature, the data from these assays could be reframed as linear features (for instance regions of high connectivity between chromosomes) and used as input to GDA. Despite the large amount of computation involved, GDA can be run on large genomes with large feature sets in ∼1 week. The most time-consuming step is repeat finding and we are exploring alternatives that would bring the overall run time down substantially.

## Methods

### Genome Decomposition Analysis pipeline

Version 1.0 of GDA was used throughout, with default parameters unless otherwise specified. A window size of 5kb was used throughout as this represents roughly the size of a gene in apicomplexans (e.g. Plasmodium spp.). The GDA v1.0 code was cloned from a private git repository to a Linux server and a Conda environment that includes all software dependencies established using the *create_gda_conda_env.py* script provided. This installation was used for running the feature extraction, clustering and analysis parts of the pipeline. A separate GDA installation and conda environment was set up on a MacBook for running the Shiny application.

The pipeline extracts the values of various sequence features (e.g. GC content) with a sliding window (default size 5kb) along all sequences in the assembly. The values are stored as separate bedgraph files (one per feature). The pipeline consists of a master script that is written in Nextflow (Di Tommaso et al. 2017). The rest of the code of the pipeline has been written mostly in Python. The Nextflow script triggers multiple third party software tools that are used to detect genomic features. As an alternative to using the Conda environment, the pipeline and its dependencies are packaged as a Singularity (Kurtzer et al. 2017) image, thus simplifying its installation in a shared environment.

Using a genome assembly FASTA file as the input, the genomic feature extraction pipeline determines low complexity sequence content using Dustmasker 1.0.0 (NCBI Resource Coordinators 2018), tandem repeat content using Tandem Repeats Finder 4.09.1 (Benson 1999), coverage of simulated reads using WGSIM 1.0 (https://github.com/lh3/wgsim), retrotransposons using LTRharvest and LTRdigest from GenomeTools 1.6.1 (Gremme et al. 2013), inverted repeats using einverted from EMBOSS 6.6.0 (Rice et al. 2000) and repeat families using either RepeatMasker + RepeatModeler 2.0.1 (Flynn et al. 2020) or Red (05/22/2015) + MeShClust2 2.3.0 (Girgis 2015; James et al. 2018). GC%, AT skew, GC skew, and the frequency of CpG, stop codons and telomeric motifs in each window are determined using Python code. If the user does not provide the pipeline with a gene annotation file, the pipeline can annotate genes itself using Augustus 3.3.3 (Stanke & Waack 2003), tRNAscan-SE 2.0.6 (Lowe & Eddy 1997), and Barrnap 0.9 [https://github.com/tseemann/barrnap]. It is possible to provide hints for Augustus using annotation transfer from a GFF3 file of a related genome with Liftoff 1.6.1 (Shumate & Salzberg 2020). With additional input data, the pipeline can detect ectopic mitochondrial and apicoplast sequences using BLAST 2.10.1 (NCBI Resource Coordinators 2018), and RNA-Seq read coverage using HISAT2 2.2.1 (Kim et al. 2015). If the user provides proteome FASTA files of species that are related to the target species, the pipeline can run OrthoMCL 1.4 (Li et al. 2003). A more detailed description of the variables can be found in Supplementary Table 1. Note that telomeric motifs, stop codons and kmers are not counted if they are broken up by a border between two windows. However, in the OrthoMCL results analysis part (when calculating the values of variables per gene in the window) a gene that is split between two windows is counted as a part of both windows.

The code for the dimensionality reduction and clustering of the data from genomic windows uses the Python UMAP (McInnes et al. 2018) and HDBSCAN (McInnes et al. 2017) libraries. The scaling of variables before running UMAP is done using MinMaxScaler from the scikit-learn package (Pedregosa et al. 2011). In the script for optimising the clustering parameters (gda_parameters.py), Silhouette score, Davies-Bouldin index and Calinski-Harabasz score are calculated for each clustering result using scikit-learn. These scores help to find the clustering settings that work the best for separating the genomic windows into distinct clusters. After determining the optimal settings for n_neighbors and minimal cluster size, the pipeline runs the final clustering. Kolmogorov-Smirnov test is used to determine whether the distribution of values of a variable in a GDA cluster is significantly different from the distribution of the values of the same variable in the rest of the genomic windows. The test is performed using the ks_2samp function from the scipy package (Virtanen et al. 2020). The Fisher test with Benjamini-Hochberg multiple hypothesis testing correction (using scipy.stats (Virtanen et al. 2020) and statsmodels.stats.multitest libraries (Seabold & Perktold) are used to determine if some types of cluster junctions occur with a different frequency than what is expected by chance. For example, this test yields a statistically significant result when windows belonging to a given cluster are located next to windows belonging to the same other cluster significantly more often than expected by chance.

While the clustering and visualisation parts of the GDA pipeline rely on bedgraph files, none of the third party software tools used by GDA produce output files in bedgraph format. We therefore use Python code written for the GDA pipeline to derive bedgraph files from the diverse set of output files produced by the third party tools. In some cases, the output of a software tool is first converted to GFF format and then the GFF file is converted to a bedgraph file. All bedgraph files corresponding to one assembly are merged into a tab-separated table. The code for merging bedgraph files into a table and for downsampling the table has been written in C++ instead of Python, in order to gain execution speed.

In this work, we distinguish four different feature sets: *seq* requires only the genome sequence as input, *gene* features are derived from a set of gene annotations, *rep* features derived from running the RepeatModeler repeat classification and analysis tool, *orth* derived from running the OrthoMCL tool for determining orthologous and paralogous relationships between protein-coding genes. These feature sets are frequently combined, as stated. In this work “full feature set” refers to the combination of these four feature sets. GDA is capable of generating additional feature sets and any arbitrary genome data tracks can be added to incorporate novel features.

### Datasets

Genome sequences and annotation for the following species were downloaded from VEuPathDB release 51 (https://toxodb.org/toxo/app/downloads/release-51/) - *Plasmodium falciparum* 3D7, *P. knowlesi* H, *P. chabaudi* AS, *P. vivax* P01, *Toxoplasma gondii* ME49, *Babesia bovis* T2Bo, *B. microti* RI, *Theileria annulata* Ankara, *T. parva* Muguga and *Cryptosporidium parvum* IowaII. Features in the GFF files labelled *protein_coding_gene* were changed to *gene*. *Eimeria tenella* Houghton data was downloaded from ENA (https://www.ebi.ac.uk/ena/browser/view/GCA_905310635.1). For OrthoMCL runs (excluding large genome analysis), all the above species were included.

### Analysis of Plasmodium falciparum

The feature extraction module of GDA was initially run using just the sequence as input, producing the following features: at_skew, cag_freq, cpg_percentage, dustmasker_low_complexity_percentage, einverted_inverted_repeat, N_percentage, gc_percentage, gc_skew, kmer_deviation_kmer_size_3, kmer_deviation_kmer_size_4, LTRdigest_protein_match, LTRdigest_LTR_retrotransposon, stop_codon_freq, tandem_repeats_fraction, telomere_freq, wgsim_depth_minimap2. A description of these features is available in Supp. Table 1.

The clustering_params function of GDA was used to determine suitable clustering parameters, with all combinations of *n neighbours* (n) = {5, 10, 15, 20} and *minimum cluster size* (c) = {50, 100, 200 500} explored. Parameter values were chosen to minimise the percentage of unclassified windows and maximise the silhouette score. This was achieved with n = 5 and c = 50. Feature extraction was also performed with the addition of gene annotations (*seq+gene*), resulting in the following additional features: exon_count, gene_average_exon_length, gene_average_intron_length, gene_length, mRNA_annotations, pseudogene_annotations, rRNA_annotations and tRNA_annotations. Clustering parameters were n = 10 and c = 40. To this feature set, repeat identification with RepeatModeler was added (*seq+gene+rep*), incorporating sum_of_simple_repeats, sum_of_complex_repeats, as well as numerous, specific simple and complex repeat family features. Clustering parameters for this feature set were n = 15, c = 50. The final feature set added features derived from an analysis of orthologues across the Apicomplexan phylum: apicomplexa_ortholog_count, apicomplexa_paralog_count, apicomplexa_protein_conservation_ratio and apicomplexa_species_specific_proteins_ratio (*seq+gene+rep+orth*). Here, the clustering parameters were chosen as n=20, c=50.

We wanted to determine whether cluster 3 (var/rif genes) and 4 (smaller multigene families) regions in the seq+gene+rep+orth run of *P. falciparum* contained genes that were more or less well conserved and more or less well covered by HP1 chromatin modifications in internal regions versus subtelomeres. To examine conservation, we looked at protein conservation ratio values for genes in cluster 3 or 4 in internal vs. subtelomeric locations. Instead of using protein conservation ratios from across apicomplexa as we had done in running GDA, we reran GDA using only predicted proteomes from *Plasmodium* species in the *Laverania* group. These were chosen specifically as they are phylogenetically close to *P. falciparum* and the *var*, *rif* and *stevor* multigene families are present, whereas they are not present outside of the group. *P. adleri* G01, *P. billcollinsi* G01, *P. blacklocki* G01, *P. reichenowi* G01 and *P. praefalciparum* G01 protein sequences were downloaded from PlasmoDB v51. We defined subtelomeric windows as those within 200kb of chromosome ends. To test whether there was a difference in HP1 occupancy between subtelomeric and internal multigene family regions, bedgraph files of log2 ratios of HP1 in trophozoites were downloaded from PlasmoDB, originally derived from (Fraschka et al. 2018). We used *bedtools intersect* to identify genes overlapping windows of each cluster. Boxplots were drawn using the *graphics* v4.0.2 package in R. Kolmogorov-Smirnov tests, using the *stats* v4.0.2 package in R, were used to determine statistical significance.

### Analysis of *P. vivax* and *P. knowlesi*

Full feature sets (*seq+gene+rep+orth*) were used for *P. vivax* and *P. knowlesi*. For *P. vivax* we chose parameters n=20, c=50, for *P. knowlesi* n=10, c=50.

### Analysis of *Eimeria tenella*

We used parameters n = 10 and c = 100 with the *seq* feature set, resulting in exclusion of 2.74% of windows and a silhouette score of 0.53. The default CAG repeat feature was excluded because this feature was originally added specifically to help identify repeats in *Eimeria* spp. Here, we wanted to demonstrate that these repetitive regions could be identified without prior knowledge. We added *rep* features (n = 5, c = 50, silhouette score = 0.32), then *gene* features (13.68% unclassified windows and silhouette score = 0.18, n=10, c=50), then *orth* features (6 clusters, n=10, c=100, 1.9% windows unclassified).

Homopolymeric Amino Acid Repeats (HAARs) were identified using Python regular expressions, looking for runs of A, S, Q, L and N of at least 7 in predicted protein sequences. There were 13,389 A, 9,404 Q, 5,350 S, 334 L and 6 N repeats.

### Analysis of *Toxoplasma gondii*

Non-chromosomal contigs were removed from the assembly. The *seq+gene+rep+orth* feature set was used with parameters n = 20, c = 50, resulting in 5 clusters, with no unannotated windows.

### Analysis of large genomes

The GDA feature extraction pipeline was run with four genomes of increasing size, with and without RepeatModeler (*rep* feature set) to show how resource requirements scale. Each was run with orthologue analysis (*orth*), genome annotation (*gene*) feature sets as well as NUclear Mitochondrial DNA (NUMT) identification. All jobs were executed on the Wellcome Sanger Institute compute farm and were given 30 Gb memory and 16 threads. Genomic windows size was 5 kb in all runs - which represents 654,762 windows for *H. sapiens*. Gene annotations were read from existing GFF files from the same origin as the assembly FASTA files (PlasmoDB, NCBI or WormBase ParaSite).

*Plasmodium falciparum* 3D7 (PlasmoDB release 43) was used with the Pf_M76611 (PlasmoDB) mitochondrial genome reference and reference proteomes *P. chabaudi chabaudi* AS, *P. ovale curtisi* GH01, *P. gallinaceum* 8A, *P. malariae* UG01, *P. berghei* ANKA, *P. vivax* P01 (from PlasmoDB release 52). *Caenorhabditis elegans* (RefSeq GCF_000002985.6) was used with mitochondrial sequence NC_001328.1 (NCBI) and predicted proteomes GCF_000001215.4 Release 6 (*Drosophila melanogaster*), GCF_000146045.2 R64 (*Saccharomyces cerevisiae*) and GCF_000001405.39 GRCh38.p13 (*Homo sapiens*) from NCBI, GCA_900184235.1 (*Oscheius tipulae*) and GCA_000469685.2 (*Haemonchus contortus*) from GenBank and PRJEA36577.WBPS14 (*Schistosoma mansoni*) from WormBase ParaSite. *Schistosoma mansoni* (WormBase ParaSite release 14, assembly Smansoni_v7) was used with mitochondrial sequence NC_002545.1 (NCBI) and predicted proteomes PRJDA72781.WBPS14 (*Clonorchis sinensis*), PRJEB527.WBPS14 (*Schistocephalus solidus*), PRJEB122.WBPS14 (*Echinococcus multilocularis*), PRJEA34885.WBPS14 (*Schistosoma japonicum*), PRJNA179522.WBPS14 (*Fasciola hepatica*), PRJEB124.WBP from WormBase ParaSite (Howe et al. 2017). Homo sapiens (NC_012920.1; NCBI) was run with mitochondrial sequence NC_012920.1 (NCBI) and predicted proteomes GCF_000002035.6_GRCz11 (*Danio rerio*), GCF_001663975.1 (*Xenopus laevis* v2), GCF_000001635.27_GRCm39 (*Mus musculus*) from NCBI.

### Data Availability

The GDA pipeline, instructions on how to install and run the pipeline and detailed results of individual analyses presented here are provided through our GitHub page: https://github.com/eeaunin/gda.

## Acknowledgements

We would like to thank Steven Doyle (Wellcome Sanger Institute) and Matthieu Muffato (WSI) for critical comments on the manuscript, Steven Doyle, Matthieu Muffato, Ksenia Krasheninnikova (WSI), Alan Tracey (WSI) and Avril Coghlan (WSI) for testing the GDA software.

