## Supplementary Material for "Characterising genome architectures using Genome Decomposition Analysis"

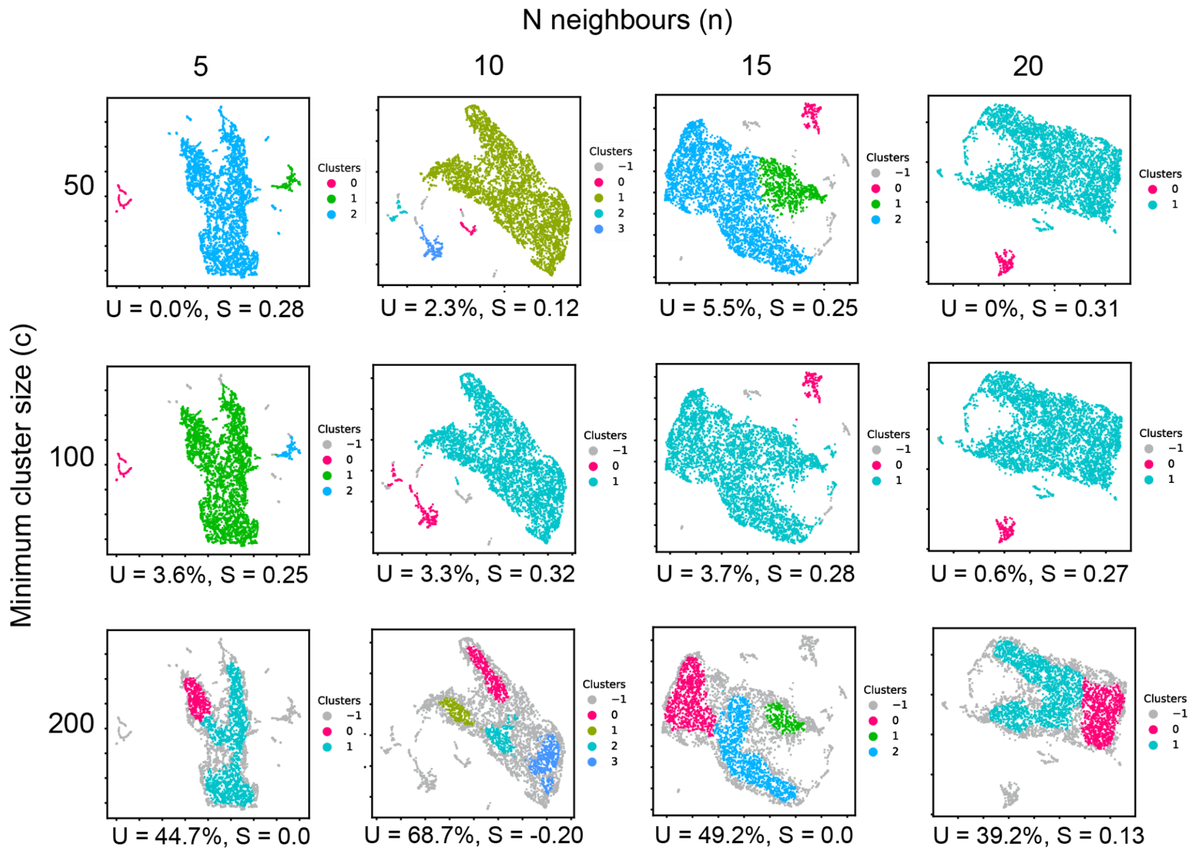


**Supplementary Figure 1. Effect of varying *N neighbours* (n) and *minimum cluster size* (c) parameters on clustering of 5kb windows from *Plasmodium falciparum* with the *seq* feature set.** Values for the percentage of unclassified windows (U) and the silhouette score (S) are shown beneath each UMAP plot. We aimed in this work to identify clustering parameters which resulted in a small percentage of unclassified windows, a high silhouette score and a reasonable number of clusters. Here we picked n = 5, c = 50, where there were no unclassified windows and the silhouette score was reasonably high. Other clusterings had higher silhouette scores (e.g. n = 20, c = 50), but had fewer clusters, suggesting they might be missing an interesting architectural feature captured by the n = 5, c = 50 clustering.

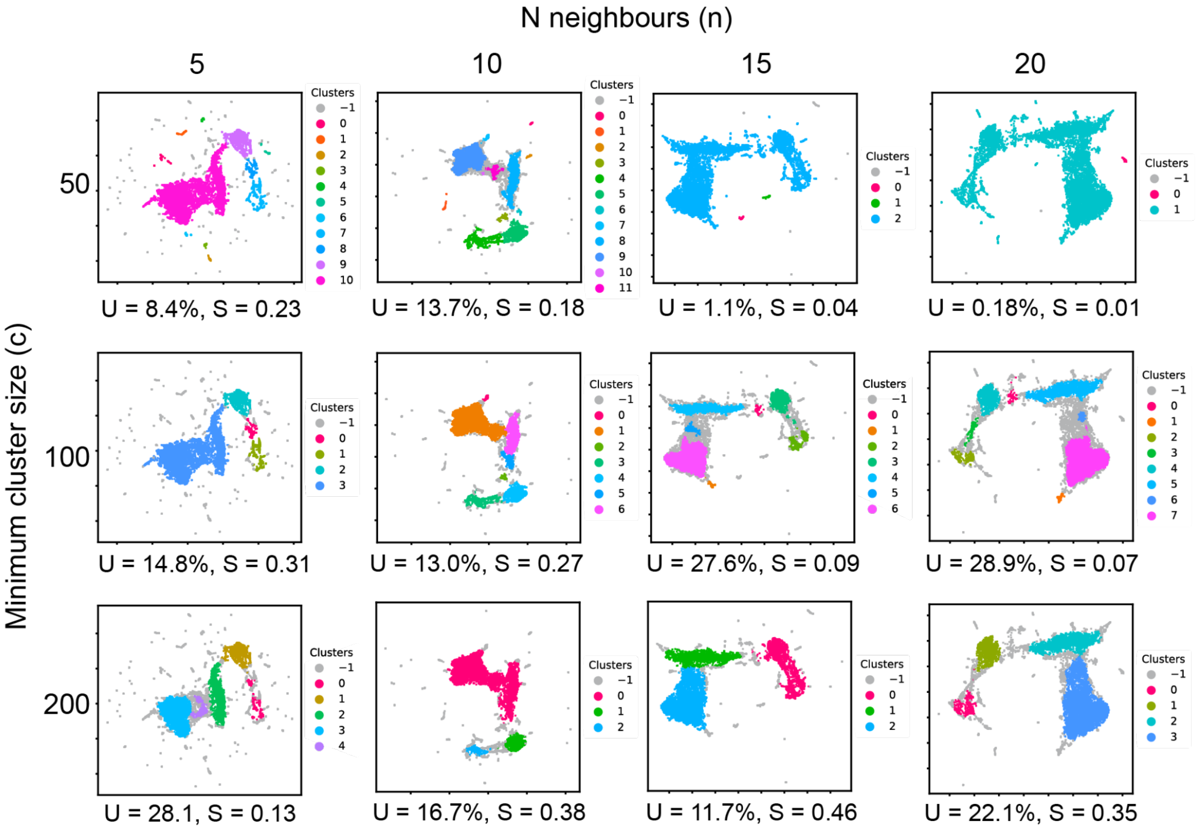


**Supplementary Figure 2. Effect of varying *N neighbours* (n) and *minimum cluster size* (c) parameters on clustering of 5kb windows from *E. tenella* with the *seq+rep+gene* feature set.** A range of *n* and *c* parameters were evaluated to determine a good clustering of genomic windows. U = unclassified window percentage, S = silhouette score. Selecting n = 10 and c = 50 allowed the identification of gene-poor subtelomeric (and sometimes internal) regions with repeat-rich regions still well-characterised.

**Supplementary Table 1. Variables extracted by the genomic feature extraction pipeline of GDA.** Each feature that can be generated by the GDA feature extraction pipeline is described here, highlighting whether it is included in a particular feature set.

| **GDA feature** | **Feature set** | **Description** |
|---|---|---|
| at_skew and gc_skew | seq | AT skew and GC skew of the sequence are calculated in Python using the standard formula for this (Perna & Kocher 1995) |
| cag_freq | seq | CAG trinucleotide frequency (this is a prominent repeat sequence in *Eimeria* (Reid et al. 2014) |
| complex_repeats_bedgraph | rep | Complex repeat families detected using RepeatMasker + RepeatModeler 2.0.1 (Flynn et al. 2020) |
| cpg_percentage | seq | CpG dinucleotide frequency (determined using string search in Python) |
| dustmasker_low_complexity_percentage | seq | Low complexity sequence percentage per window. It is determined by running Dustmasker 1.0.0 (NCBI Resource Coordinators 2018) with default settings and counting masked nucleotides per window |
| ectopic_mitochondrion and ectopic_apicoplast | - | Putative ectopic mitochondrion or apicoplast sequences, detected using BLAST against user-provided mitochondrial or apicoplast sequence. Nucleotide BLAST of the target genome (query) against a reference organellar sequence |

| | | |
|---|---|---|
| | | (subject) is run with E-value cutoff 1e-30. If the length of BLAST hit alignment is 90% or more of the length of the query sequence, it is assumed that the query sequence is probably a real rather than an ectopic organellar sequence. Sequences with shorter alignment lengths than that are reported as putative ectopic organellar sequences |
| einverted | seq | Inverted repeats, detected by running einverted from EMBOSS 6.6.0 (Rice et al. 2000) with default settings. The output of einverted is reformatted as a GFF and a bedgraph file using Python |
| gene_length, exon_count, gene_average_exon_length, gene_average_intron_length | gene | Features determined based on mRNA gene annotations. Average mRNA gene length, average exon count per mRNA gene, average exon length and average intron length. The values of these variables are calculated per gene in the window |
| gene_dna_strand_bias | - | Gene DNA strand bias shows the tendency of mRNA genes to be all on the same strand in the window. The value is 1 if all genes in the window are on the same strand (it does not matter which one). The value is 0 if genes in the window are equally distributed between both strands.<br>This is an optional feature and is not calculated by default |
| gaps | seq | Assembly gaps (Ns). Windows containing assembly gaps are excluded from UMAP+HDBSCAN clustering |

| | | |
|---|---|---|
| gc_percentage | seq | DNA GC% (calculated using Python) |
| kmer_deviation | seq | kmer skew for a particular kmer length (how much the distribution of kmers in the window differs from what is expected by change, given the GC content of the sequence in the window).<br>The probabilities of a randomly selected base in the sequence being A, T, G or C are calculated based on the GC content of the sequence. E.g. if the GC content is 60%, the probabilities of a randomly selected nucleotide in the sequence or its reverse complement being A, T, G or C are the following. A: 0.2, T: 0.2, G: 0.3, C: 0.3.<br>For finding the probability of observing a kmer, the probabilities of all individual nucleotides in the kmer are multiplied. When counting kmers in the sequence, the counts of each kmer are aggregated with the counts of their reverse complement kmers. To account for this, the probability of observing the kmer is multiplied by 2 (so that the probability value describes the chance of a kmer or its reverse complement kmer).<br>The expected count of a kmer is found by multiplying the previously calculated probability of observing the kmer with the total number of observed kmers. For each kmer in the window, the absolute value of the difference between observed and expected counts is recorded. kmer deviation for the window is the sum of these differences for all kmers in the window. |

| | | |
|---|---|---|
| ltrdigest_protein_matches | seq | Matches to retrotransposon proteins detected by LTRdigest from GenomeTools 1.6.1 (Gremme et al. 2013). LTRdigest performs a search for a set of HMM (hidden Markov model) files. This set contains a selection of HMMs from PFAM (Sonnhammer et al. 1997) that have been described as associated with LTR retrotransposons by Steinbiss et al. (Steinbiss et al. 2009). It also contains the HMMs from The Gypsy Database (GyDB) (Llorens et al. 2011), downloaded in 2020 |
| ltrdigest_retrotransposons | seq | Putative retrotransposons (detected using LTRharvest and LTRdigest from GenomeTools 1.6.1 (Gremme et al. 2013). Only the sequences containing LTRdigest protein matches are counted |
| mRNA_annotations | gene | mRNA gene density (either from user-provided gene annotations or detected from de novo annotations using Augustus 3.3.3 (Stanke & Waack 2003)).<br>Augustus is run with default settings and requires the user to select the Augustus species model. In order to make Augustus run faster, the target genome is split into chunks and Augustus is run in parallel with each of the chunks, using GNU parallel version 20201122 (Tange 2011). It is possible to provide hints for Augustus using annotation transfer from a related genome with Liftoff 1.6.1 (Shumate & Salzberg 2020) with default settings |

| | | |
|---|---|---|
| pseudogene_annotations | gene | Density of pseudogenes. Pseudogene features are read from a user-provided GFF file, if they are present there. The pipeline does not annotate pseudogenes *de novo* |
| rRNA_annotations | gene | Density of rRNA genes. rRNA gene annotations are either read from a user-provided GFF file or created *de novo* using Barrnap 0.9 (https://github.com/tseemann/barrnap) with default settings. The user is required to select a value for the kingdom parameter of Barrnap |
| tRNA_annotations | gene | Density of tRNA genes. tRNA gene annotations are either read from a user-provided GFF file or created *de novo* using tRNAscan-SE 2.0.6 (Lowe & Eddy 1997) with default settings |
| simple_repeats_bedgraph | rep | Simple repeat families detected using RepeatModeler+RepeatMasker 2.0.1 (Flynn et al. 2020). Using a Python script written specifically for the GDA pipeline, the sequences have been collapsed to count repeats that are the reverse complement of one another as the same repeat. The sequences have also been collapsed to count the repeats that are identical if the starting point is adjusted as the same repeat (e.g. TGGTT is the same as GGTTT) |
| ortholog_count | orth | Average number of orthologs in other species for genes within a window. Diamond 2.0.4 blastp (Buchfink et al. 2021) is used to produce the BLAST results that will be used as an input for OrthoMCL 1.4 (Li et al. 2003). |

| | | |
|---|---|---|
| paralog_count | orth | Average number of paralogs within the same species averaged across genes in the window |
| protein_conservation_ratio | orth | The proportion of species that have orthologs for each gene, averaged across the window. The value is 1 if all genes in the window have orthologs in all other species in the OrthoMCL run. The value is 0 if none of the target species proteins in the window have any orthologs in any other species. |
| HISAT2_RNA-Seq_coverage | - | RNA-Seq read FASTQ files are validated using ValidateFastq 0.1.1 [https://github.com/biopet/validatefastq]. RNA-Seq reads are mapped to the target genome using HISAT2 2.2.1 (Kim et al. 2015). The coverage of the mapped RNA-Seq reads per genomic window is found using SAMtools (Li et al. 2009). |
| species_specific_proteins_ratio | orth | The number of genes with no OrthoMCL orthologs in the window divided by the number of all genes in the window. The value is 1 if none of the target species proteins in the window have orthologs, and 0 if all of the target species proteins in the window have orthologs. |
| stop_codon_freq | seq | Stop codon frequency, based on the counts of TAG, TAA and TGA trinucleotides in the DNA sequence |
| sum_of_complex_repeats | rep | Sum of values of RepeatModeler + RepeatMasker tracks for complex repeat families |

| | | |
|---|---|---|
| sum_of_simple_repeats | rep | Sum of values of RepeatModeler tracks for simple repeat families |
| tandem_repeat_density | seq | Density of tandem repeats detected using Tandem Repeats Finder 4.09.1 (Benson 1999). The settings used for Tandem Repeats Finder are the following:<br>Match: 2, Mismatch: 1000, Delta: 1000, PM: 80, PI: 10, Minscore: 25, MaxPeriod: 1000, -m, -h, -ngs. The same settings have previously been used in (Reid et al. 2014). |
| telomere_freq | seq | Telomeric sequence frequency. It is determined by finding exact matches to one or multiple telomeric sequence motifs that are selected in the run settings with the --telomeric_seq_preset option (defaults to vertebrates, i.e. the TTAGG motif). The telomeric sequence presets are based on sequences from the Telomerase Database (Podlevsky et al. 2008). All matches to the telomeric motif are counted, including those that are not located at scaffold ends |
| wgsim_minimap2_coverage | seq | This variable is used to estimate mappability. To find the values for this, simulated Illumina reads are derived from the target genome using WGSIM 1.0 (https://github.com/lh3/wgsim). By default, the number of reads that is generated is such that it should yield 10x coverage. The read lengths are 150 bp and the random seed is set to 1. The rest of the settings for WGSIM are left as default. The simulated reads are mapped to the target |

| | | |
|---|---|---|
| | | genome using Minimap2 (Li 2018) with short read mapping settings ("-ax sr"). Using SAMtools, the BAM file with mapped reads is filtered to keep only reads with mapping quality 60. A Python script is then used to convert coverage values into bedgraph format |
